# Preclinical shortwave infrared tumor screening and resection via pHLIP ICG under ambient lighting conditions

**DOI:** 10.1101/2022.09.07.506973

**Authors:** Benedict Edward Mc Larney, Mijin Kim, Sheryl Roberts, Magdalena Skubal, Hsiao-Ting Hsu, Anuja Ogirala, NagaVaraKishore Pillarsetty, Daniel Heller, Jason S. Lewis, Jan Grimm

## Abstract

There is a critical need to improve optical imaging that will lead to its widespread acceptance for routine clinical procedures. Shortwave infrared (SWIR, 900–1700nm) imaging has demonstrated clear advantages over visible and near-infrared imaging (reduced autofluorescence with improved contrast, resolution, and sensitivity at tissue depth). Here we show that the previously reported compound, pH low insertion peptide (pHLIP) conjugated to indocyanine green (ICG, pHLIP ICG) currently in clinical trials, serves as an excellent candidate for SWIR imaging protocols. SWIR’s increased sensitivity enabled preclinical tumor screening and resection at exposure times as low as 0.1 ms with acceptable signal-to-noise and contrast-to-noise ratios. Imaging was performed under ambient lighting conditions, and SWIRs sensitivity enabled an extended surgical resection window up to 96 hrs post injection in an orthotopic breast cancer mouse model. This work provides a direct precedent for the clinical translation of SWIR pHLIP ICG imaging for cancer resection.

**One Sentence Summary:** SWIR imaging under ambient lighting is highly sensitive to pHLIP ICG, a cancer targeting fluorescent agent currently under clinical investigation.

## INTRODUCTION

The advent of efficient fluorescent proteins and dyes in combination with single photon sensitive camera devices has delivered a plethora of pre- and clinical implementations of optical imaging for cancer resection.(*1, 2*) These optical agents and devices commonly perform imaging in the visible (VIS, 400–700 nm) and near infrared (NIR, 700-900 nm) light spectrum.(*1, 3-5*) Biophotonic imaging below 650 nm presents significant drawbacks in terms of endogenous tissue absorption and scattering, ultimately reducing the effective resolution, imaging penetration depth and is susceptible to high levels of tissue autofluorescence from laser excitation.(*6, 7*) To overcome these limitations, there have been significant efforts by various groups to facilitate imaging above 650 nm where tissue optical properties are more favorable.(*1, 8-11*) Indocyanine green, one of the few clinically approved fluorescent dyes, takes advantage of the decreased tissue optical aberrations by imaging at NIR wavelengths above 800 nm and has found a variety of applications.(*12-16*) Whilst the perturbations imposed by tissue are reduced at these wavelengths, the silicon-based detectors used to perform imaging have rapidly reduced sensitivity above 800 nm (quantum efficiency of 60% or lower).(*17, 18*) Silicon detectors currently provide a reliable platform and are widely employed, but have rapidly decreasing quantum efficiency above 800 nm where ICG is most fluorescent.(*19, 20*)

Shortwave infrared (SWIR, 900-1700 nm) imaging is poised to revolutionize pre- and clinical biophotonic imaging.(*18, 21, 22*) SWIR detectors are comprised of Indium Gallium Arsenide (InGaAs) chips with a deposited indium phosphide layer preventing detection of light below ∼920 nm and extending to ∼1700 nm.(*18, 23*) SWIR detectors typically have a high (above 80%) quantum efficiency within this range, and by thinning of the indium phosphide gap can have improved sensitivity below ∼920 nm.(*24-26*) SWIR detectors originally found use in military, industrial and scientific applications where they provided increased contrast and resolution when imaging through visible light scattering environments e.g., fog, astronomy or food inspection.(*27-29*) In the SWIR spectral range of light, tissue scattering, absorption and autofluorescence are of negligible levels resulting in improved contrast, higher penetration depths and improved resolution at depth.(*6, 7, 30-32*) Additionally, due to the reduced absorption of tissue at these wavelengths it is safe to input more laser excitation energy when compared to VIS and NIR wavelengths.(*32, 33*) The combination of these factors enables background free, high sensitivity imaging with an abundance of optical photons that remain in the ballistic regime over extended distances when compared to visible imaging.(*32, 34*) Due to the spectral response of human eyes (380-720 nm) and the insensitivity of the SWIR detectors below 920 nm, it is feasible to perform SWIR imaging without the need for ambient light removal at no effect to human vision under suitable lighting conditions.(*35, 36*) We believe this is critical for widespread adoption as it requires minimum modification to existing facilities with both surgeons and patients being more comfortable if procedures can be performed under ambient light conditions.

SWIR imaging is often unknown in preclinical imaging settings, largely due to the unavailability of imaging systems, dyes, and agents for routine applications to investigate biological systems that are complicated and diverse. This availability is further exacerbated in clinical imaging where system requirements are stricter than preclinical imaging. Recently, SWIR imaging has characterized the extended emission of ICG from 900–1500 nm (not detected by silicon-based sensors) with notable improvements in sensitivity, resolution and contrast compared to silicon-based imaging.(*7, 20, 31, 37, 38*) The superiority of SWIR imaging in a preclinical setting was demonstrated using porcine angiography model where the improved sensitivity enabled blood vessel detection beneath blood pools, not visible on widely used silicon detectors.(*39*) In addition to ICG imaging there has been a rapid development of dedicated SWIR fluorescence agents for in vivo applications.(*40, 41*) One area which would greatly benefit from SWIR imaging is that of clinical cancer resection. In comparison to functional imaging of e.g., calcium dynamics, cancer resection provides a more facile translation for SWIR imaging where imaging can be performed with a binary output of tumor/no tumor for resection if a dye can be targeted and localized to a tumor. The current array of preclinically validated SWIR targeting agents will still require further validation and toxicology studies before reaching clinical deployment.(*36, 40, 42*) Clinical SWIR translation has predominantly focused on non-targeted ICG based imaging and has already shown significant improvements over NIR imaging for liver tumor surgery, glioma resection, cystic renal mass removal and brain metastasis.(*43-46*)

Fortunately, pHLIP ICG (pH low insertion peptide, conjugated to ICG), which until this point has not been validated with SWIR imaging, is a tumor targeting agent that is currently undergoing clinical trials for imaging-guided breast cancer resection (NCT05130801).(*47-50*) In acidic pH environments, typical of tumor microenvironments, the pHLIP compound will insert into cellular membranes, displaying high selectivity and contrast over healthy tissues and organs.(*47*) Whilst the current clinical trial and this work is focused on breast cancer, pHLIP ICG has also been shown to preclinically delineate a variety of cancers in mouse models.(*47*) Additionally, pHLIP can be conjugated to other optical dyes e.g., QC1 for optoacoustic imaging and can be radiolabeled for PET imaging.(*51, 52*) In this work, we combine the sensitivity of SWIR imaging to ICG with the tumor selectivity of pHLIP for preclinical tumor resection in an orthotopic murine breast model. We show that a commercially available SWIR imaging system has significantly improved sensitivity over widely used commercial imaging system IVIS^®^ for ICG imaging.(*47, 53-55*) The improved sensitivity enables extension of the surgical resection window from 24 to up to 96 hrs. We also show increased contrast of the tumor at 72 hrs over non-cancerous tissue, performed tumor screening and resection confirmation with exposure times ranging from 10 to 0.1 ms at all time points and most notably show that SWIR pHLIP ICG imaging could be carried out under ambient lighting conditions, significantly enhancing clinical practicality and translation. This work provides a direct step for the clinical translation of SWIR pHLIP ICG imaging for cancer resection, which could further aid the clinical translation of other SWIR cancer targeting agents.

## RESULTS

### SWIR pHLIP ICG ambient light imaging setup and characterization

The first step of these experiments was to establish the capability of SWIR imaging under ambient lighting conditions without detrimental effects on image quality or sensitivity. The finalized system setup is shown along with image processing workflow is shown in Figure 1A. We employed an open-source Arduino kit to power a programmable red, green, and blue (RGB, white) light LED to provide ambient lighting within the preclinical SWIR imaging enclosure.

**Fig 1.**
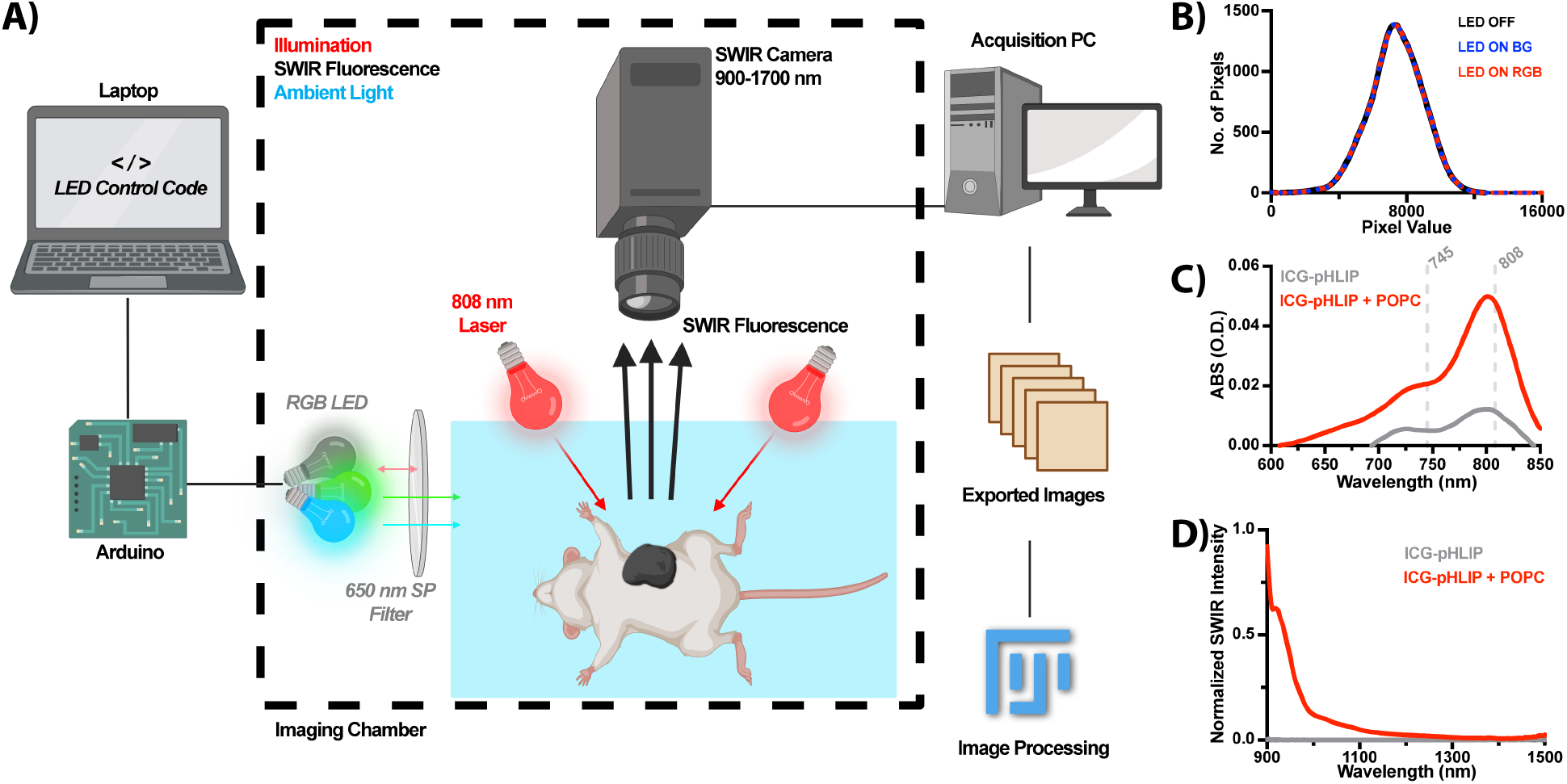
Schematic of the SWIR pHLIP ICG imaging setup. **A)** A dedicated preclinical SWIR imager was adapted for ambient light imaging of pHLIP ICG. An LED controlled via an Arduino provided illumination like that of room lighting (<650 nm) within the enclosure. SWIR pHLIP ICG fluorescence was excited with an 808 nm laser and detected using an InGaAs sensor. Recorded images were then converted to tiff files in Fiji (ImageJ) where automated scripts performed image correction and analysis. **B)** Histogram response of the SWIR sensor without LED illumination, with the LED in a blue, green (BG) state and with the LED in a red, green, and blue (RGB) state. No significant difference was found between these states highlighting the detectors insensitivity to ambient lighting under these imaging conditions. **C)** The absorption profile of 100 μL of an 8 μM pHLIP ICG solution with (fluorescent) and without (essentially non-fluorescent) the presence of POPC-liposomes. The excitation wavelengths of NIR (745 nm) and SWIR (808 nm) systems are shown. **D)** The extended SWIR emission of pHLIP ICG in both low (no POPC) and highly (with POPC) fluorescent states.

The system required setup in this manner to avoid SWIR contamination from non-LED room lighting outside the enclosure and for safe laser containment during imaging. All Arduino and LED power was provided via a laptop USB port. The system sensitivity was tested with LED states in full power for RGB (white) and GB (cyan) combinations. The LED was housed in a custom 3D printed enclosure coated with aluminum foil (for light reflection) and a 650 nm short pass filter mounted in front of the LED. The resulting histogram distribution of pixel values acquired under high sensitivity and without laser excitation is shown in Figure 1B (see also Supplemental Figure 1). No difference could be determined between LED Off, LED GB and LED RGB modes with subtraction of the LED Off frames from LED GB and RGB states showing no discernable signal. This allowed us to conclude the insensitivity of the system to ambient lighting within the enclosure under these conditions. Following confirmation of the systems insensitivity to ambient lighting, we measured the NIR absorbance and SWIR emission spectra of pHLIP ICG on respective dedicated spectrometers. As shown in Figure 1C, 100 μL of an 8 μM pHLIP ICG solution displayed a characteristic ICG absorption band, peaking at 802 nm in the presence of POPC-liposomes. The liposome free solution displayed reduced absorption due to the lack of membrane insertion, as expected. In figure 1D, we show that the fluorescence mechanism (consistent with the previously reported NIR imaging) of pHLIP ICG is preserved in the SWIR region and extends to approximately 1500 nm.(*47, 48*)

### Advantage of SWIR imaging for pHLIP ICG

Preclinical mouse optical imaging is often carried out on an IVIS®. The IVIS has capabilities of imaging multiple (5) mice at a single time with a wide range of excitation, emission, and luminescent options. The versatility and ease of use of this system has resulted in multiple investigations where the IVIS plays a central role e.g., tumor growth assessment via luciferase. The system employs a halogen lamp for excitation and a silicon-based sensor cooled to -90°C for detection and is capable of imaging ICG with relevant excitation and emission settings (745nm/845nm). Whilst convenient, we found the system to be relatively insensitive to ICG in comparison to the commercial SWIR system used here. We directly compared both systems in terms of sensitivity to pHLIP ICG when imaging 100 μL of an 8 μM pHLIP ICG solution in PBS in the presence of POPC liposomes with and without scattering tissue (5 mm of raw chicken breast). Our results indicate that the SWIR system was 100x more sensitive than the IVIS system with successful signal at exposure times as low as 0.1 ms (Figure 2A). This increase in sensitivity is consistent when imaging through 5 mm of tissue as shown in Figure 2B. It should be noted there are some slight caveats with this comparison. Both systems performed imaging without binning (the SWIR sensor is incapable of on-sensor binning, unlike the IVIS) for easier comparison. Additionally, the IVIS halogen emission spectrum is very weak past 700 nm (745 nm excitation used here) with additionally reduced sensitivity of the sensor past 800 nm whilst the SWIR system incorporates a high power 808 nm laser for excitation (pHLIP ICG’s peak absorption).

**Fig 2.**
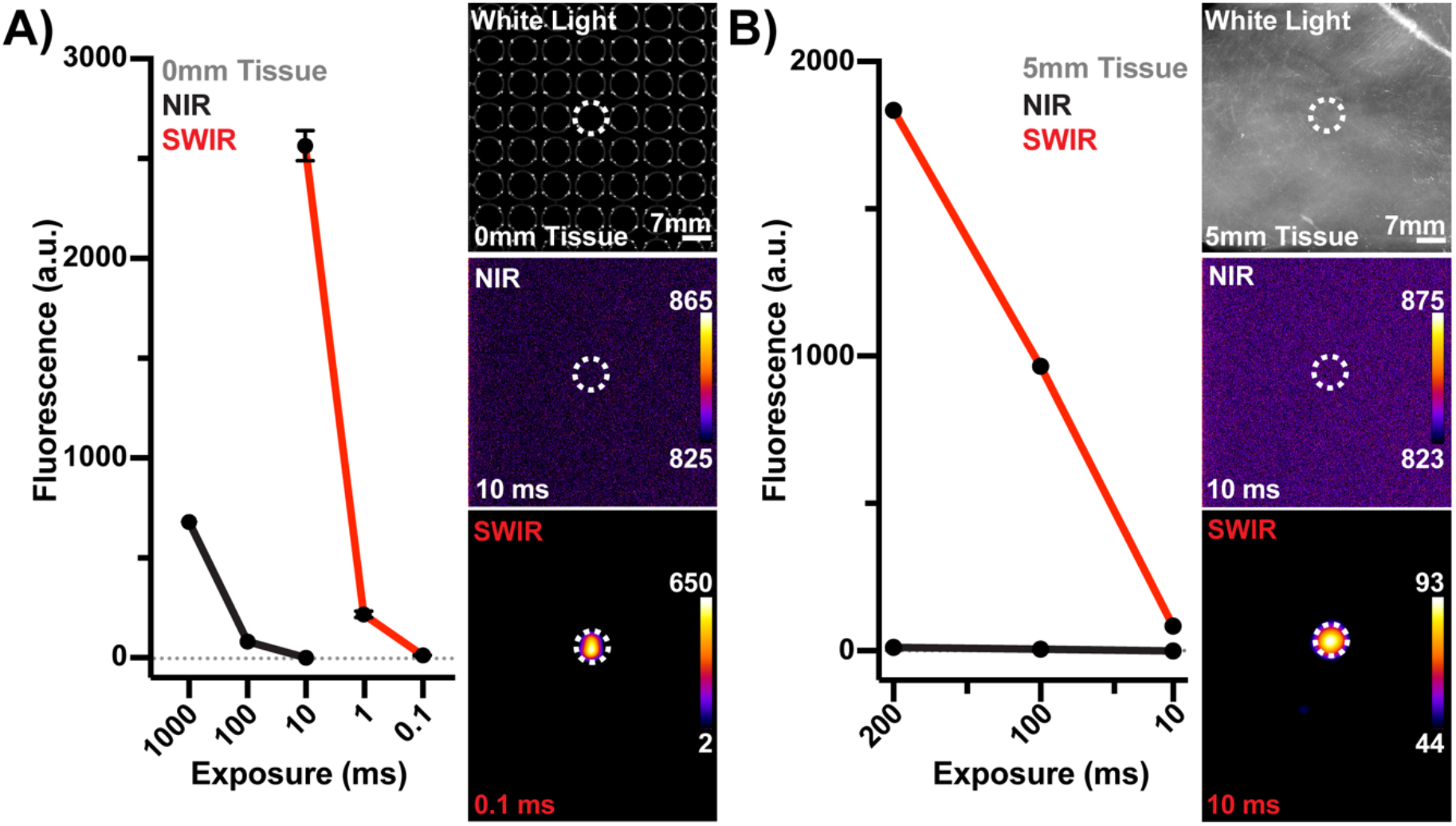
Phantom comparison of commercial NIR and SWIR imaging. **A)** Left, quantification of the reported mean gray values of both the commercial NIR (IVIS, 745 Exc, 840 Em) and SWIR (808 nm Exc, 900 – 1700 nm Em) systems at exposure times from 1 – 0.1 ms. Images are taken from a 96 well plate with a single well containing 100 μL of an 8 μM pHLIP ICG solution in PBS containing 10% POPC-liposomes. Right, representative images of both modalities highlighting SWIRs improved sensitivity. **B)** Left, quantification of the reported mean gray values of both modalities when imaging through 5 mm of scattering tissue (raw chicken breast). Right, representative images of both modalities with SWIR capably imaging through tissue at 10 ms exposure times.

### Extended pHLIP ICG imaging timeline and sensitivity

A previous publication reporting pHLIP ICG for the delineation of tumors provided NIR fluorescence levels for the biodistribution of pHLIP ICG for numerous mice at timepoints up to 48 hrs.47 We analyzed this data highlighting that tumor and liver fluorescence were in direct competition and found that post 48 hrs, the tumor contrast should be significantly improved, see Supplemental Figure 2. However, detecting this contrast improvement would only be feasible assuming the employed imaging system could provide sufficient sensitivity. To confirm this sensitivity to pHLIP ICG, we imaged various concentrations of the agent at differing exposure times, see Supplemental Figure 3. With laser exposure well within ANSI limits (103.3 mW/cm2 of a 330 mW/cm2 limit at 808 nm), we found the threshold of SWIR imaging for pHLIP ICG to be in the singular nanomolar range (4 nM) at exposure times of 10 ms. This would potentially correspond to an imaging rate of approximately 100 Hz aside from the camera being unable to transfer images at those rates under these sensitivity settings (∼30 Hz). As expected, the sensitivity decreased one log as exposure time decreased one log highlighting the systems linear sensitivity to pHLIP ICG concentration (10 ms: 4 nM, 1 ms: 40 nM and 0.1 ms: 400 nM).

### SWIR pHLIP ICG tumor screening under ambient lighting

Having determined the SWIR system provided suitable sensitivity, notably improved to that of the IVIS, we then investigated the potential to detect increased tumor contrast up to 96 hrs post pHLIP ICG injection. Athymic female nude mice (hairless, FoxN1^nu^) were utilized to enable facile SWIR visualization of deep-seated organs through the skin without hair or pigmentation hampering detection. Mice were spontaneously injected with murine breast cancer cells (4T1) into the mammary fat pad located closest to the liver and allowed to proliferate for 7–9 days.(*47*) Once tumors had reached a suitable size four mice were intravenously injected with pHLIP ICG at 0.5 mg/kg and imaged at 1, 24, 48, 72 and 96 hrs post injection with one mouse receiving no injection. Due to the increased depth of the liver compared to the orthotopic breast tumor model location, we performed imaging slightly above ANSI limits (808nm, 450 mW/cm^2^). This ensured potential signal collection for the deep-seated liver without sacrificing mice at extended imaging timepoints. In figure 3A, we show the tumor screening potential of SWIR pHLIP ICG imaging at 1, 24, 48, 72 and 96 hrs post injection. SWIR white light images were captured via illumination with a 1300 nm LED with the ambient lighting LED clearly shown in the bottom right corner of all imaging points. In some timepoints and orientations, reflections from the mouse fluorescence can be seen. SWIR fluorescence images are shown for all injected mice (0.5 mg/kg) at all timepoints. As expected, the liver shows the highest fluorescence levels at 1 hr post injection and decreases at later time points, with the tumor increasingly standing out. We assessed the SNR of the system at all timepoints under the exposure times tested as shown in Figure 1B. The system consistently performed with high SNR values above a minimum acceptable threshold of 5 dB. Following this, we measured regions of interest from the tumor and surrounding area to track the clearance rates of pHLIP ICG. The retained fluorescence of the tumor is prominent past 24 hrs in comparison to surrounding areas. Finally, we calculated the CNR of pHLIP ICG SWIR imaging comparing tumor contrast to the surrounding body. The 1-hour timepoint is not shown as this was a negative CNR value due to high liver uptake. We found that CNR increased linearly, peaking at 72 hrs and slightly decreasing at 96 hrs. Exposure times of 10 and 1 ms provided similar CNR values at all timepoints. To ensure the potential for clinical translation of these results imaging was carried out on a second batch of mice within ANSI limits (808nm, 300 mW/cm^2^) with similar results shown (see Supplemental Figure 7).

**Fig 3.**
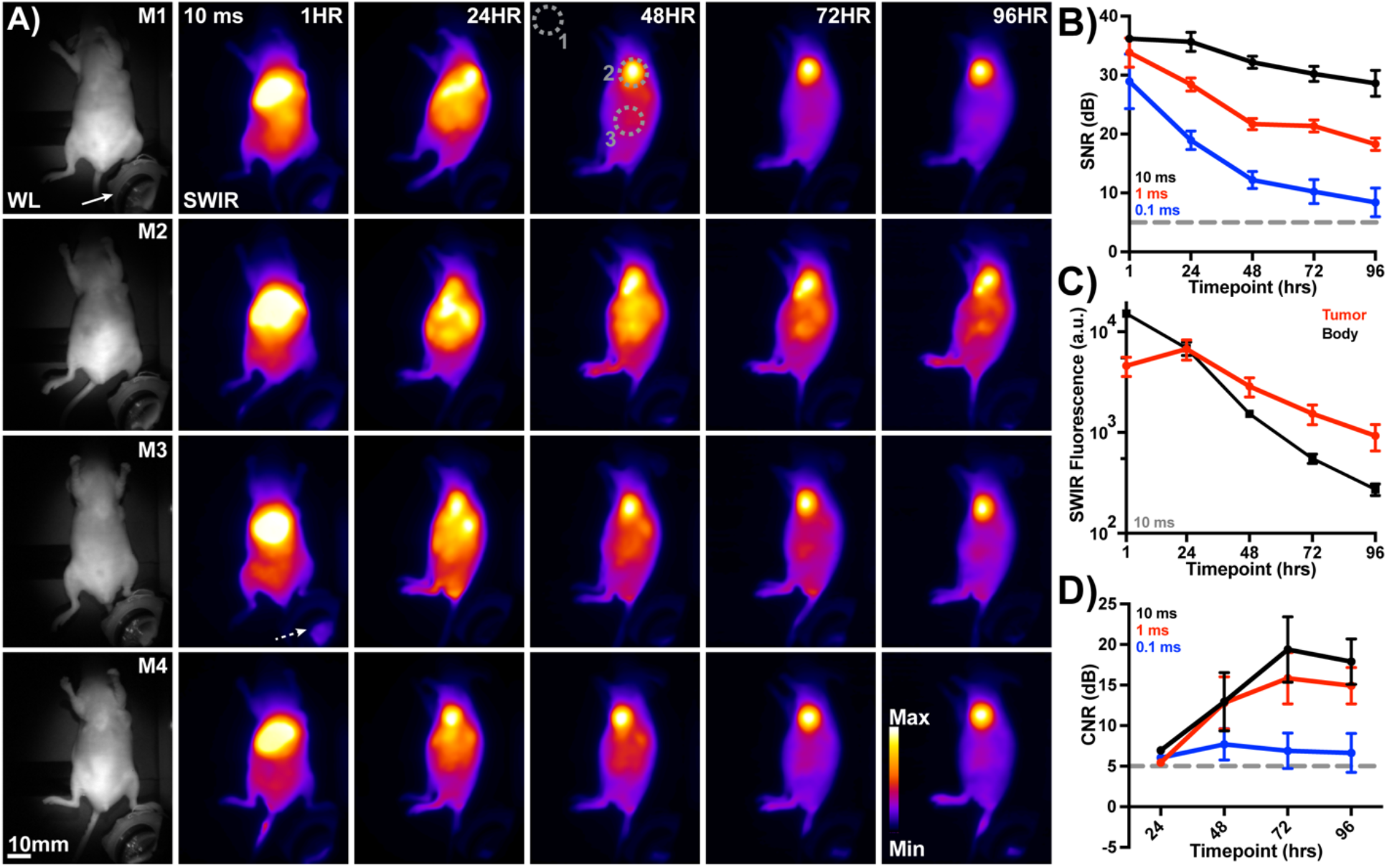
SWIR based tumor screening with pHLIP ICG under ambient lighting. **A)** White light (WL, 1300 nm LED illumination) SWIR images of pHLIP ICG injected mice (0.5 mg/kg) at 1 hr post injection. The solid arrow indicates the ambient light source (LED). SWIR fluorescence images of injected mice (n = 4) at 10 ms exposure times under 808 nm excitation at 1, 24, 48, 72 and 96 hrs post pHLIP ICG injection. Images are thresholded to respective min and max values with mice positioned to show brightest fluorescence locations e.g., the liver at 1 hr and tumor at 48 hrs (ROI 2). The dotted arrow highlights a fluorescence reflection in the image. Regions of interest (ROIs) are shown by the dotted gray lines where 1 represents system noise, 2 the tumor values and 3 the body values. Negligible signal was detected from a control mouse, data not shown (non-injected, n=1). **B)** The signal to noise ratio (SNR) of the brightest point in dB from all mice from 1 to 96 hrs. A threshold of 5 dB was chosen for image acceptability. C) Comparative SWIR fluorescence values (10 ms exposure) from all mice comparing tumor and body fluorescence levels. Tumor fluorescence is retained up to 96 hrs with brighter fluorescence past 24 hrs. D) Contrast to noise ratios (CNR) in dB of the tumor to body SWIR fluorescence highlighting the increase in contrast past 24 hrs at exposure times of 10, 1 and 0.1 ms. Data from n = 4 biological replicates are shown in all cases aside from D, 24 hrs where only n = 1 value is shown for all. Mean and standard deviation are shown in all cases.

### SWIR pHLIP ICG tumor resection under ambient lighting

Having shown that SWIR pHLIP ICG screening was feasible up to 96 hrs post injection, we then performed surgical resection (post euthanasia) on these mice. In the four mice tested the tumor was readily delineated from non-tumor tissue (see Figure 4A). In these cases, both the SNR and CNR were of acceptable levels (above 5 dB, see Figure 4B). Finally, to ensure the accurate distribution of pHLIP ICG, biodistribution studies were performed on all mice. In all cases the tumor showed a higher fluorescence level than other tissues, readily differentiated and confirming the enhanced clearance of pHLIP ICG from tissues other than the tumor. Representative images are shown in Figure 5A, with fluorescence values for all organs from all mice shown in Figure 5B.

**Fig 4.**
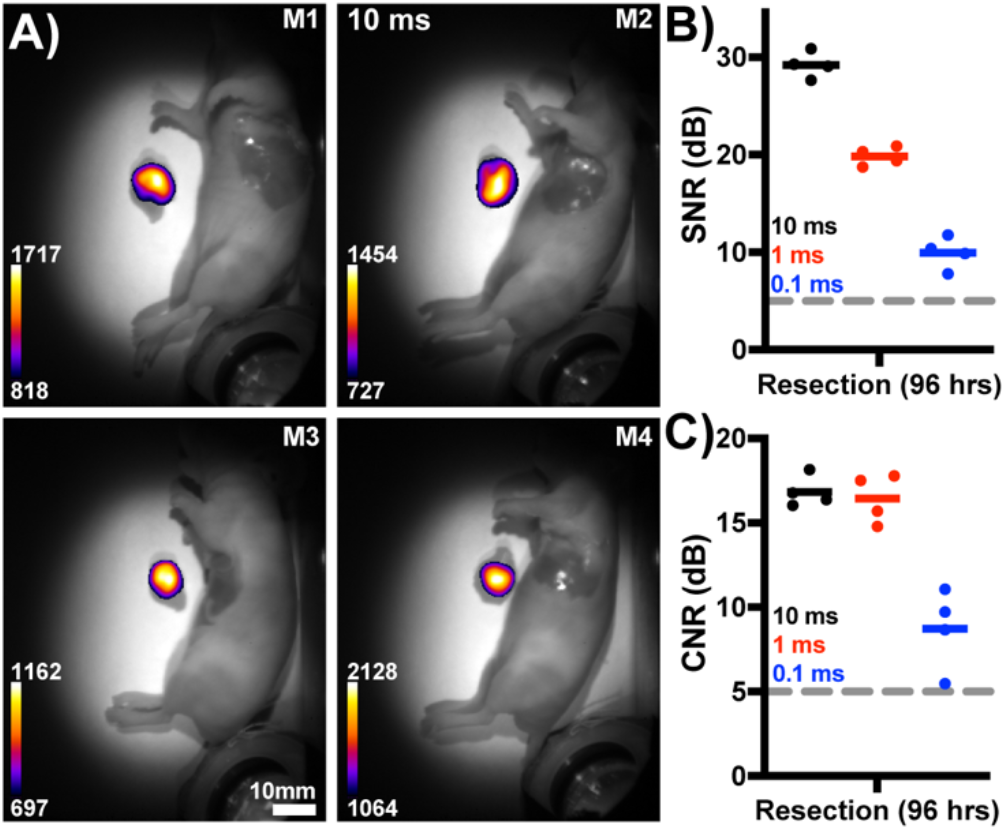
pHLIP ICG SWIR guided resection under ambient lighting. **A)** WL overlaid SWIR fluorescence images of post euthanasia resection via pHLIP ICG, 96 hrs post injection. In all cases the tumor is clearly delineated from the body. Mice are the same as those in Figure 3. **B)** Quantified SNR of the resected tumors at all exposure times. **C)** Quantified CNR levels of the tumor compared to body fluorescence for all exposure times. Acceptable levels are defined by the 5 dB level in both graphs. Data for n = 4 biological replicates is shown in all cases along with the mean.

**Fig 5.**
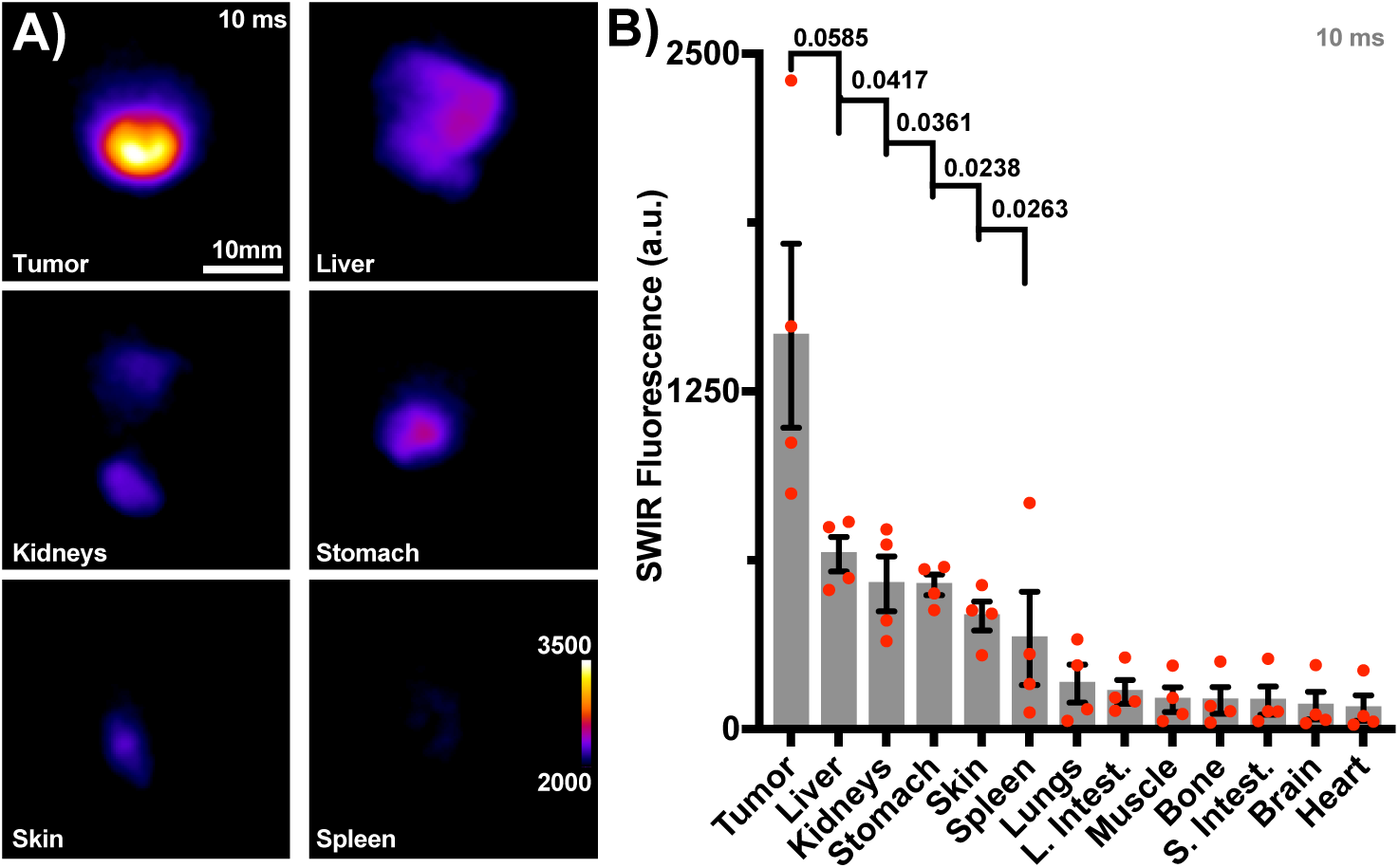
SWIR necropsy biodistribution at 96 hrs under ambient lighting. **A)** Representative SWIR pHLIP ICG images of the tumor, liver, kidneys, stomach, skin, and spleen post biodistribution. Images are thresholded to show signal from other organs. **B)** Decreasing SWIR fluorescence values of all organs from all mice. Endogenous signal values from a non-injected control mouse, have been respectively subtracted for each tissue type. The mean (gray bar), SEM (black bars) and individual values (n = 4, red dots) are shown in all cases. P values comparing the tumor to other tissues are shown for those in A). Organs are from mice used in Figure 3 and 4.

## DISCUSSION

In the preclinical space there has been a significant effort to highlight the advantage of SWIR imaging over conventional NIR imaging, development of novel probes and the development of improved imaging setups.(*21, 31, 39, 41, 56*) This work is the first to show that a pH based tumor targeting agent currently undergoing clinical trials is also suitable for SWIR imaging. We have shown that the pHLIP peptide when conjugated to ICG fluorescence retains it NIR fluorescent properties in the SWIR region, where tissue scattering, absorbance, and autofluorescence are minimal (see Figure 1).(*47*) Whilst the number of photons emitted by pHLIP ICG diminish past 850 nm, the available photons are readily detected with minimal background interference reminiscent of Raman spectroscopy. When comparing two commercially available preclinical systems, the advantages of SWIR imaging are even more obvious. In this investigation SWIR clearly outperformed the IVIS system achieving SNR and CNR values of 33.9 and 12.9 dB at 10ms exposure times at 96 hrs post injection whilst IVIS detected no signal at 10 ms and required 10 s exposure times to achieve similar respective SNR and CNR values of 19.3 and 19.5 dB (see Figure 2 and Supplemental Figures 7 and 8). Certain caveats should be noted with this comparison. Firstly, the IVIS system employs a halogen lamp source with weak emission in the ICG excitation band. Secondly, the silicon-based sensor present in the IVIS has decaying quantum efficiency (QE) past 800 nm. In comparison, the SWIR sensor used here is not capable of on-sensor binning like that of the IVIS, employs a less efficient lens (SWIR f# of 1.4 versus the IVIS f# of 1.0) and additionally can only image one mouse at a time reducing throughput. However, the SWIR system is capable of laser excitation at the absorption peak of pHLIP ICG, see Figure 1. These direct factors could be overcome in custom designs but are outside the scope of this work comparing commercially available solutions.

Aside from these technological differences, previous work has already established the superiority of SWIR imaging over conventional methods.(*31, 39*) We briefly reanalyzed the clearance rate of pHLIP ICG from a previous publication to optimize the tumor to body contrast ratios.(*47*) This analysis revealed that the liver was responsible for the majority of pHLIP ICG uptake, with the tumor and liver reaching similar fluorescence levels approximately 48 hrs post injection. Based on the peak uptake timepoints, clearance rates for both the tumor (14 hrs post injection) and liver (4 hours post injection) were extrapolated predicting enhanced tumor contrast imaging past 48 hrs (see Supplemental Figure 2). However, imaging at these extended timepoints required suitable sensitivity, which was readily achieved with the SWIR system (4 nM sensitivity at 10 ms exposure times with 808 nm excitation well within ANSI limits, 103.3 mW/cm^2^, see Supplemental Figure 3).(*33*)

Having assessed the advantage of pHLIP ICG SWIR imaging in phantoms, we then investigated its utility in tumor screening and resection. The SWIR system and accompanying image processing pipeline produced relevant and accurate images at all timepoints from all mice (see Figure 3 and Supplemental Figure 4). Video rate SWIR imaging was achieved with acceptable SNR and CNR values at exposure times of 10, 1 and 0.1 ms at all timepoints (see Supplemental Video 1). We preformed surgical resection and biodistribution assessments of SWIR’s capabilities at 96 hrs post injection (see Figures 4, 5 and Supplemental Figure 5). SWIR pHLIP ICG tumor uptake was confirmed by H&E staining (see Supplemental Figure 6). The enhanced sensitivity of SWIR imaging facilitated video rate tumor detection, guided resection and biodistribution at relatively low probe levels and as late as 96 hrs post injection with high SNR and CNR levels, and median necropsy-based muscle to tumor ratios of 13.38 at 96 hrs. Most notably all the SWIR images and results presented here were performed under ambient lighting conditions with no detriment to signal accuracy.(*41, 42*) Considering the wide tumor targeting capabilities of pHLIP ICG, other tumors should also be investigated preclinically for SWIR imaging.(*47*) Furthermore, we have shown the potential for the direct clinical translation of these results by detecting similar trends, SNR and CNR values at reduced laser intensities within ANSI limits (300 mW/cm^2^, see Supplemental Figure 7).

The results presented here have a direct effect on preclinical and clinical tumor imaging. Future preclinically tested targeting agents (peptides, nanoparticles, small molecules, antibodies etc.) investigations can be readily assessed via SWIR imaging at video rates by conjugating ICG. From a pre- and clinical perspective, the enhanced sensitivity of SWIR imaging may enable a reduction in the required dose for detection. This may ultimately enable picomolar sensitivity and aid in removing one of the current barriers for the clinical translation of fluorescent agents. We have also shown that this sensitivity is feasible with ambient lighting conditions but would require LED installation as opposed to the current phosphorescent bulbs used in most settings (which contain a SWIR component in their spectrum). The sensitivity of SWIR enabled sensitivity at exposure times of 0.1 ms but surgical resection and tumor detection do not require such temporal resolutions. Whilst not possible with this system, the exposure times used here could be employed but with imaging rates of 30 Hz (30 ms). Ensuring laser triggering in combination with these shortened exposure times would reduce the overall exposure of the agent and thus reduce photobleaching effects.(*57-60*) A knock-on of this would be an extended surgical window and surgical imaging time.

This work has focused on the SWIR detection of ICG conjugated pHLIP, however this peptide could be amenable to other dyes which have shown further improved SWIR applicability.(*40, 41*) For example, pHLIP conjugation to a dye that is excited at 1064 nm where lasers are widely available would enable an approximately three times increase in the total light exposure with further reduced optical aberrations from tissue.(*6, 33, 36*) This will enable further improved detection of deeply seated tumors for screening, or remnant tumor tissue during resection.(*39*) The clinical applicability of pHLIP ICG now calls for a clinically appropriate SWIR system and resection setup that can translate these results to directly improve both patient welfare and clinical tools.

## MATERIALS AND METHODS

### Study Design

The primary goal of this study was to establish the suitability of the cancer targeting fluorescent dye, pHLIP ICG, for SWIR imaging. SWIR imaging has already shown significant advantages for ICG detection over NIR and pHLIP ICG could readily avail of these advantages.(*20, 31*) The majority of SWIR clinical imaging is performed without targeted dyes and preclinical SWIR agents require lengthy validation before clinical trial commencement.(*40, 43*) A clinically relevant tumor targeting dye would have direct impacts on translating SWIR imaging to the clinic and could ultimately improve successful surgical resection. To test our hypothesis, we firstly assessed the SWIR spectrum of pHLIP ICG on a custom-built SWIR spectrometer and validated that the NIR mechanism of pHLIP ICG functioned in the SWIR regime. Following this we reanalyzed the previously published NIR pHLIP ICG data to show via a semilog extracted decay function that tumor contrast would increase and surpass other organs 48hrs after injection assuming sufficient sensitivity.(*47*) We ensured this sensitivity by imaging pHLIP ICG solutions in its fluorescent form ultimately achieving 4 nM sensitivity at 10 ms exposures and at 103.3 mW/cm^2^ 808 nm laser excitation (within ANSI limits).(*33*) We then preclinically tested our method by injecting four 4T1 orthotopic tumor bearing mice with 0.5 mg/kg of pHLIP ICG imaged them along with a single control every 24 hours from 1 – 96 hrs post injection at varying exposure times. Imaging was performed slightly above ANSI limits to ensure liver detection, the main competitor to the tumor for pHLIP ICG contrast. A second batch (n = 5) mice were imaged in the same manner but with reduced laser exposure (under ANSI limits) for SWIR imaging and direct comparison to the current gold standard IVIS imaging. In all cases tumor resection and organ necropsy was performed at 96 hrs post injection for all mice, confirming the hypothesis. No blinding, randomization or sample size calculations were performed. Data analysis and statistical components have been outlined where used including schematic diagrams, identified regions of interest, calculation formulae and all software used. No data was excluded from the analysis. All mouse handling, imaging, and housing was performed in accordance with NIH guidelines and under approved IACUC protocols at MSKCC.

### SWIR Spectral Measurement of pHLIP ICG

The fluorescence emission spectra of pHLIP ICG was assessed in the presence of and without POPC (1-Palmitoyl-2-oleoyl-sn-glycero-3-phosphocholine) liposomes (100nm in size, T&T Scientific Corp, TN, USA). pHLIP ICG has previously shown a high NIR fluorescent state when bound to liposomes and low NIR fluorescent state without the presence of liposomes enabling *in vitro* testing.(*47, 48*) SWIR spectra were acquired using a home-built SWIR fluorescence spectroscopy system consisting of a tunable white light source, inverted microscope, and 1D InGaAs NIR detector. The SuperK EXTREME supercontinuum white-light laser source (NKT Photonics) was used with a VARIA variable bandpass filter accessory, capable of tuning the output 500−825 nm, set to a bandwidth of 20 nm centered at 575 nm. The light path was shaped and fed into the back of an inverted IX-71 microscope (Olympus), where it passed through a 20× SWIR objective (Olympus) and illuminated the samples in a 96-well clear flat bottom UV-transparent microplate (Corning). Emission from the pHLIP ICG (8 μM) was collected through the objective and passed through a dichroic mirror (875 nm cutoff, Semrock). The excitation wavelength was 808 nm. The light was f/# matched to the spectrometer using several lenses and injected into a Shamrock 303i spectrograph (Andor, Oxford Instruments) with a slit width of 100 μm, which dispersed the emission using a 86 g/mm grating with 1.35 μm blaze wavelength. The spectral range was 723−1694 nm with a resolution of 1.89 nm. The light was collected by an iDus 1.7 μm InGaAs (Andor, Oxford Instruments) with an exposure time of 0.1–5 seconds. An HL-3-CAL-EXT halogen calibration light source (Ocean Optics) was used to correct for wavelength-dependent features in the emission intensity arising from the spectrometer, detector, and other optics. A Hg/Ne pencil-style calibration lamp (Newport) was used to calibrate the spectrometer wavelength. Background subtraction was conducted using a well in a 96-well plate filled with PBS. Following acquisition, the data was processed with custom codes written in MATLAB that applied the aforementioned spectral corrections and background subtraction and fitted the fluorescence emission peaks with Lorentzian functions. Absorbance was measured from a fresh sample of the same solution on an Odyssey plate reader (Odyssey CLx, LI-COR).

### SWIR in vivo fluorescence Imaging

In vivo fluorescence imaging of pHLIP ICG was performed using a preclinical SWIR hyperspectral mouse imaging system (IR-VIVO & PHySpec, Photon Etc., Canada). Excitation was provided by two continuous wave 808 nm diode lasers each with an output power of 2 W. Lasers were reflected off optical mirrors and distributed over the entire mouse with a power density of 90–450 mW/cm2. Imaging at 808 nm was performed slightly above ANSI limits (330 mW/cm2 for 808 nm) to ensure signal detection from the deep-seated liver at extended time points. We have also included data at 100 mW/cm2, far below ANSI limits showing the sensitivity down to 4 nM in phantoms along with imaging a second batch of mice at 300 mW/cm2 with similar results. The camera was set to -70.0°C high gain sensor mode, 0 gain analogue to digital conversion, 8 MHz readout rate, no flat field correction and 14-point bit depth. At these settings the camera achieved data transfer rates of 30 Hz with exposure times adjusted between 10–0.1 ms. Video images were captured using the camera control software. SWIR white light images were acquired via 1300 nm LED illumination at 10 ms exposure times. Ambient lighting conditions were achieved within the enclosure via an RGB LED housed in a custom 3D printed holder and a 650 nm short pass filter (Thorlabs, FESH0650, NJ, USA) providing 70 μW/cm2 at 488 nm (room lighting outside the enclosure measured to be 140 μW/cm2 at 488 nm). The LED was powered and controlled via an Arduino and suitable software with red, green, or blue hues set to values between 0 (off) – 255 (brightest), see Supplemental Figure 1. Imaging was carried out with the red set to 0, green and blue to 255 providing cyan illumination across the FOV. The ambient lighting LED was positioned to always be visible within the FOV whilst imaging.

### IVIS Imaging

NIR images were acquired on a commercially available preclinical imager (IVIS Spectrum CT, PerkinElmer), currently a gold standard for preclinical optical imaging. The system was set to low binning, f# of 1, high lamp, 745 nm excitation and 820 nm emission filter sets. The raw (luminescent) tiff files were used for analysis post reference noise subtraction with median and gaussian blur steps applied in the same manner as SWIR processing. Images were acquired at exposure times ranging from 0.01 – 10 s.

### Image Processing

All data from the SWIR device was saved in a .h5 format. An automated ImageJ macro was used to import the image files and output them in a 14 bit .tiff format for streamlined analysis.56 A second macro for an automated image analysis pipeline consisted of the following steps: darknoise reference images were acquired for all conditions and subtracted from experimental data. After noise subtraction, a median filter (outlier removal, kernel size of 1, threshold 500) was applied followed by gaussian blurring (sigma of 2 pixels). Finally, the “Fire” LUT was applied to all fluorescence images along with thresholding. Quantification images was performed on single frames with each exposure time. The image processing pipeline is summarized in Supplementary Figure 4.

### Data Analysis

Data analysis was performed in GraphPad Prism (version 9.3.1, GraphPad Software, San Diego, California USA) and MATLAB (2021b, Mathworks, USA). Signal to noise (SNR) and contrast to noise (CNR) values (in dB) were calculated in Excel (Version 16.57, Microsoft) according to equations 1 and 2.

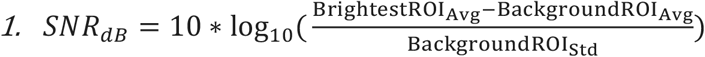

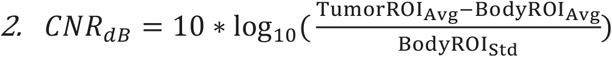

### Mouse handling

All mouse handling, imaging, and housing was performed in accordance with NIH guidelines and under approved IACUC protocols at MSKCC. Athymic nude female mice (n = 10, The Jackson Laboratory, Foxn1nu, 002019, inbred) were housed under a 12 hr on 12 hr off light cycle, 5 mice per cage with food (Sulfatrim addition) and water ad libitum. Mouse breast tumor models were generated via the injection of 0.3×106 4T1 cells (ATCC, CRL-2539 validated via STR) suspended in 30 μL of Matrigel into the mammary fat pad. 4T1 cells were validated to be mycoplasma free. Tumors were allowed to proliferate until they reached a size of ∼100 mm3 (7 – 9 days post injection) at which point mice were intravenously injected with 0.5 mg/kg of pHLIP ICG. Mice were then imaged at 1, 24, 48, 72 and 96 hrs post injection of pHLIP ICG for tumor delineation. Anesthesia was achieved in all cases via gaseous inhalation of isoflurane (3% v/v for induction, 1-2% v/v for maintenance during imaging). At 96 hrs mice were euthanized via CO2 inhalation in accordance with approved protocols. Tumors were resected with successful tumor excision confirmed via SWIR imaging. Finally, necropsy biodistribution was performed on the mice with the following organs being harvested and imaged on the SWIR imager: tumor, liver, kidneys, spleen, stomach, large and small intestines, brain, skin, bone, muscle, heart, and lungs. The respective SWIR organ values from a non-injected (control) mouse (n=1) were subtracted from injected mice values (n = 4). A second batch of mice (n = 5) were also injected with pHLIP ICG (n = 4, n = 1 control) and imaged at reduced laser exposure and for comparison with IVIS imaging. In total n = 10 mice were used for this investigation.

### Hematoxylin and Eosin Staining and imaging

Hematoxylin and Eosin (H&E) staining was performed on 5 μm thick slices using an automated system (Leica Autostainer, ST5010) with the protocol as described in Supplemental Table 1. Stained slices were then automatically imaged on a Pannoramic Scanner (3DHistech, Budapest, Hungary) using a 20X/0.8NA objective. The produced images were interpreted in SlideViewer (Version 2.5, 3DHistech) and representative regions of interest (ROIs) were chosen at 20x magnification, see Supplemental Figure 6.

## List of Supplementary Materials

Supplementary Materials File containing Fig. S1 to S8

Table S1 describing H&E staining contained in the Supplementary Material File

Movie S1 showing video rate acquisition of tumor bearing mouse injected with pHLIP ICG

## Author contributions

BML: concept, experimental design and setup, imaging parameters, data collection, image processing and analysis. MK, SR, MS: experimental design and setup, data collection, analysis. All Authors: Provided input, assisted in experimentation, writing and approval of the text.

## Data and materials availability

All data, code, and materials used in the analysis is available upon request from the corresponding authors.

