## Supplemental Material for "Preclinical shortwave infrared tumor screening and resection via pHLIP ICG under ambient lighting conditions"

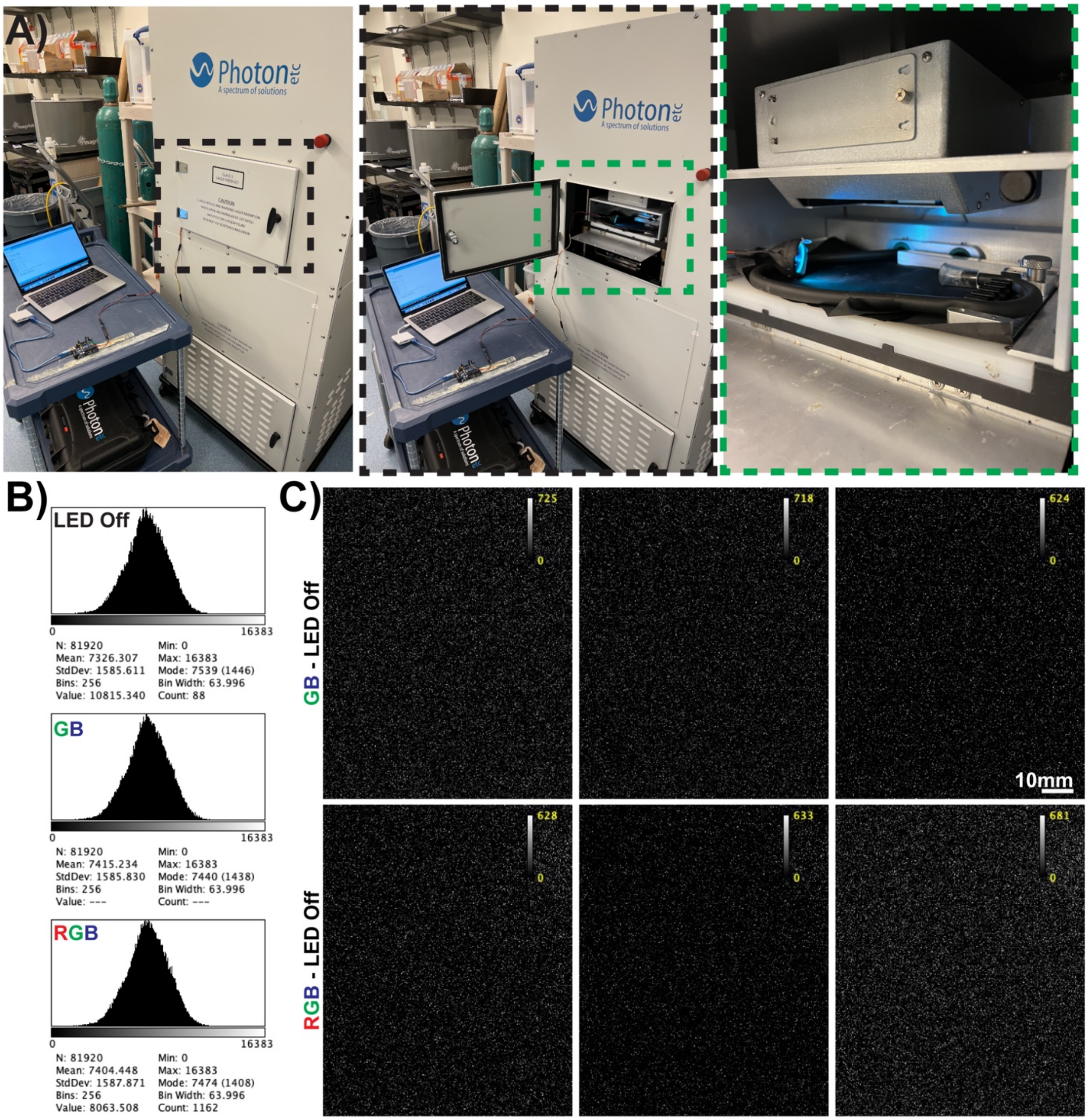


**Supplemental Figure 1. Assessing the SWIR systems sensitivity to ambient light. A)** Left, the Arduino and PC setup to provide ambient lighting inside the enclosure. Middle, the open door of the enclosure highlighting the LED location. Right, zoomed in view of the ambient lighting setup and LED lighting levels achieved in the enclosure. **B)** Histogram representation of recorded pixel values with the LED off, LED with green and blue (GB) light to max and with red, green, and blue (RGB) light set to max. **C)** Images of LED light settings post LED off frames subtraction. No light from the LED is detected by the setup, exposure time of 0.01s.

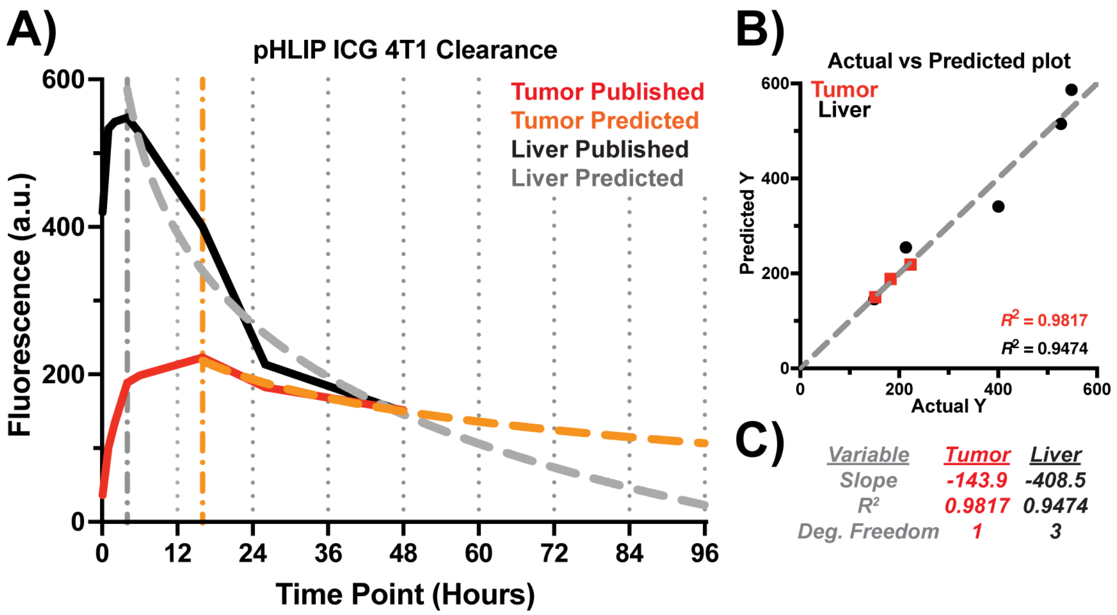


**Supplemental Figure 2. Known and predicted pHLIP ICG clearance rates up to 96 hrs post injection. A)** Graphical representation of the pHLIP ICG clearance rates for 4T1 tumors in BALB/C mice of previously published data.[46] The main competitor for pHLIP ICG uptake over the tumor (red line, published) is the liver (black line, published). The clearance rates to 96 hrs were estimated by the fitting of a semilog non-linear function from the peak signal levels (liver at 4 hours and tumor at 14 hours). The liver clearance rate is shown by the gray dotted line and tumor is the orange dotted line. **B)** The actual versus predicted plot for both tumor and liver values with respective $R^2$ values of 0.9817 and 0.9474. **C)** The fit components of the semi log non-linear fit function of A).

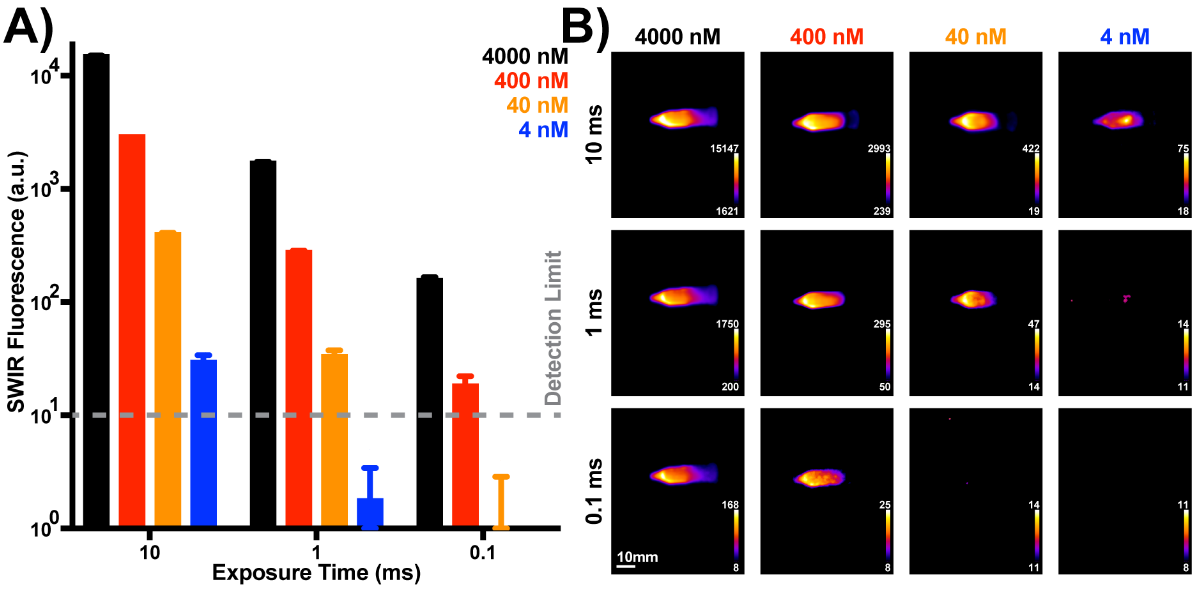


**Supplemental Figure 3. Assessing the pHLIP ICG sensitivity limits of SWIR Imaging. A)** Graphical representation of SWIR pHLIP ICG sensitivity for the exposure times and logarithmically decreasing concentrations from 4000 – 4 nM. **B)** Representative images for each exposure time and solution concentration. Images were acquired from a 1.5 mL Eppendorf containing 900 µL of a 10% POPC liposome PBS solution and pHLIP ICG at the listed concentrations. Values have been corrected via subtraction of a pHLIP ICG free solution. SWIR showed sensitivity down to 4 nM at 10 ms exposure times with 808 nm excitation at 103.3 mW/cm$^2$.

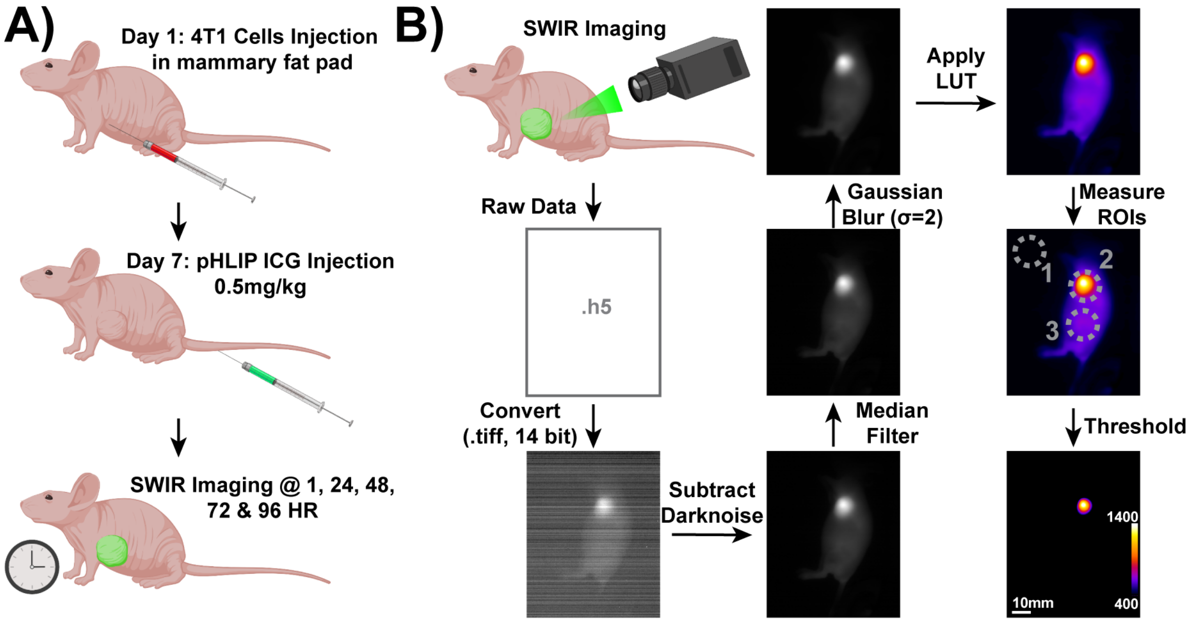


**Supplemental Figure 4. Schematic of SWIR pHLIP ICG imaging timeline and processing steps. A)** The experimentation timeline of tumor generation and pHLIP ICG imaging points. Nude mice were injected with 0.3 x $10^6$ 4T1 cells in 30 µL of Matrigel into the mammary pad. Approximately 7 days post tumor injection (tumors reached a size of 100 $mm^3$) mice received an intravenous (IV) tail injection of 0.5mg/kg of pHLIP ICG in PBS. Mice were imaged at 1, 24-, 48-, 72- and 96-hours post injection. **B)** The image processing pipeline for SWIR pHLIP ICG detection. Images were stored in .h5 formats and batch converted to 14 bitt tiff files. Following tiff conversion the images underwent appropriate dark noise reference subtraction, median filtering (1x1 kernal, 500 threshold) for outlier removal, gaussian blur application (sigma of 2 pixels) and finally look up table (LUT, Fire) application. Regions of interest (ROIs) were selected for measurements before thresholding and overlaying where necessary on corresponding white light images.

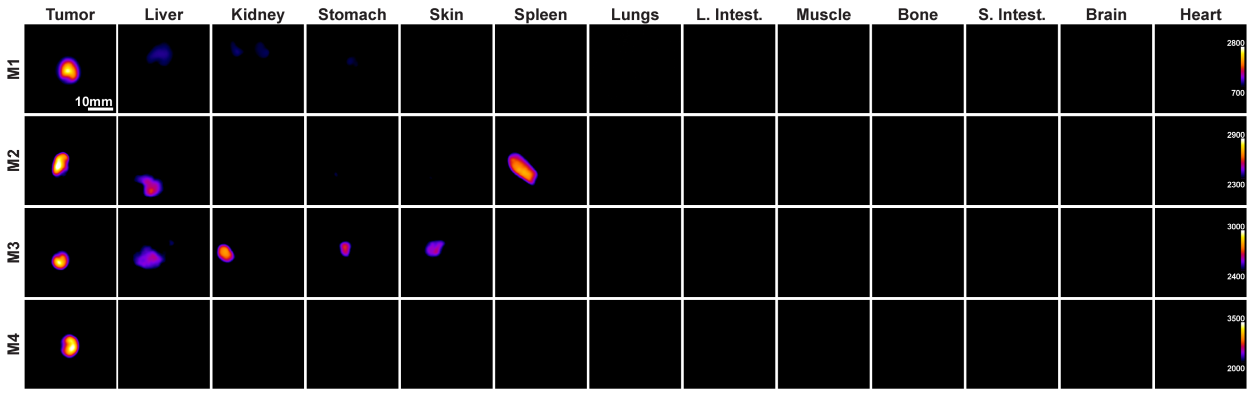


**Supplemental Figure 5. SWIR biodistribution assay of all organs from all mice.** The images of organs from all mice in the main text are shown with corresponding threshold levels. Thresholds have been applied to remove any signal not from the tumor. Images were also acquired under ambient lighting conditions and have been cropped to better show the organ location.

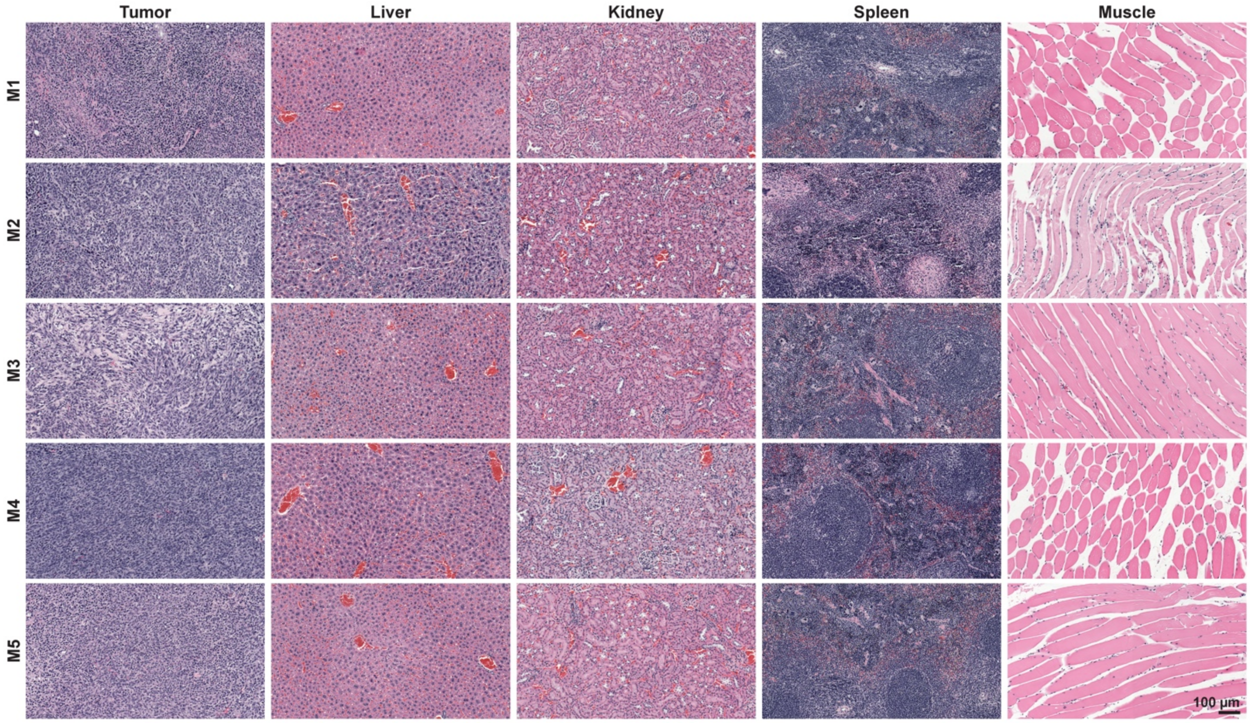


**Supplemental Figure 6. H&E staining of tumors and select organs from all mice.** M1 – M4 were injected with ICG-pHLIP (0.5mg/kg), M5 received no injection and was imaged at all timepoints under identical conditions. H&E sections are similar across all mice and all images are shown under 20x magnification.

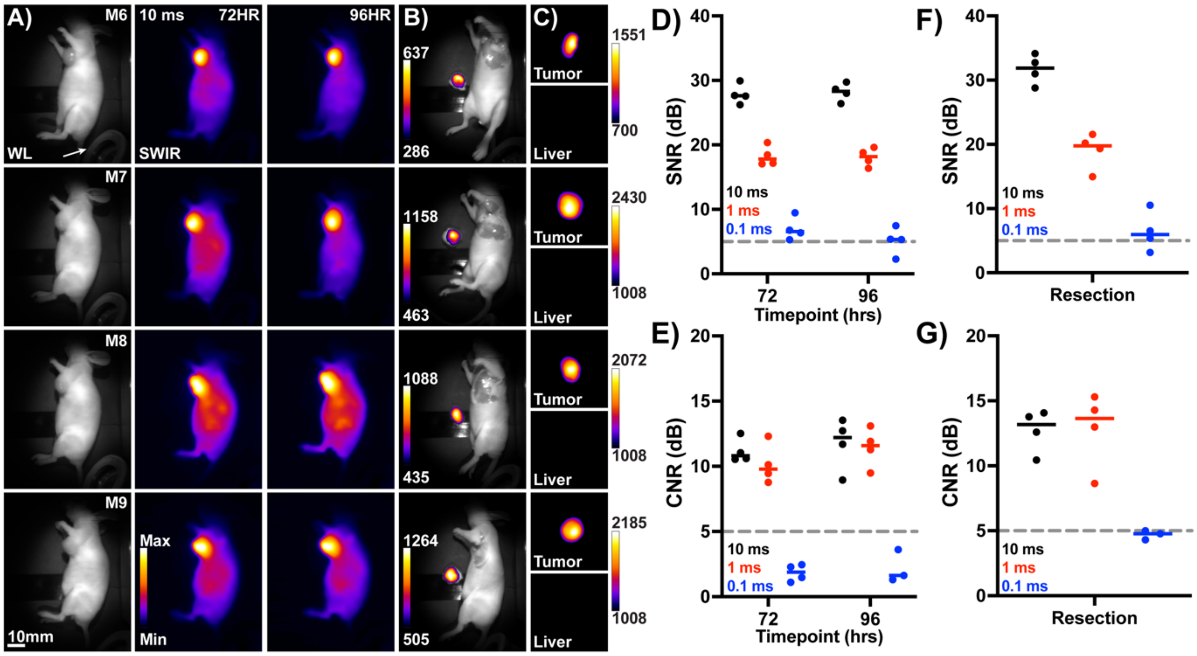


**Supplemental Figure 7. SWIR ICG-pHLIP imaging within ANSI limit excitation intensities (300 mW/cm$^2$). A)** White light (1300 nm) and SWIR fluorescence images of a second batch of mice at 72 and 96 hrs post injection of ICG-pHLIP (0.5 mg/kg). **B)** SWIR resection of mice (post euthanasia) shown in A) 96hrs post injection. **C)** Tumor and liver SWIR fluorescence levels for all mice at 96hrs. **D)** SWIR screening SNR values (dB) for all mice and all exposure times at 72 & 96 hrs for images shown in A). **E)** SWIR CNR values at 72 and 96 hrs for images shown in A). **F)** Resection SNR values for all mice and **G)** resection CNR values for all mice for images shown in B). The reduced laser intensity (300 mW/cm$^2$) prevents satisfactory detection levels at 0.1 ms for resection SNR, 72 & 96 hrs screening CNR and resection CNR.

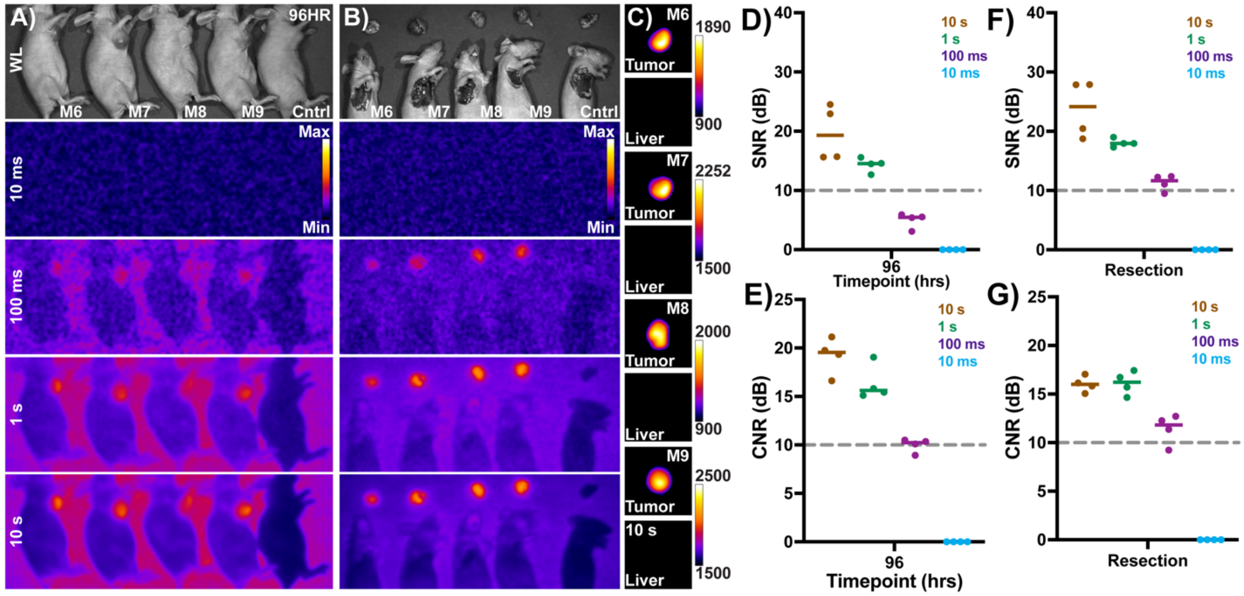


**Supplemental Figure 8. IVIS imaging of mice at 96hrs post injection for comparison to SWIR imaging. A)** ICG-pHLIP screening of mice as shown in Supplemental Figure 7 at 96 hrs post injection. IVIS settings had various exposure times as shown, 745 nm excitation, 820 nm emission, lamp high, F stop 1, small binning. **B)** Resection of mice under IVIS imaging. **C)** Confirmation of tumor and liver IVIS fluorescence levels. **D)** IVIS SNR values for all mice and exposure times. The mean SNR value at a 10s exposure time is 19.31 in comparison to the SWIR 0.01s SNR of 33.91. **E)** CNR values for mice as shown in A). **F-G)** SNR and CNR values for IVIS guided resection. In all cases the exposure time of 0.01s for IVIS imaging does not detect signal with only 1s exposure times providing the most reliable signal, 100 times slower than the SWIR imaging (0.01s).

| Step | Station | Reagent / Station Name | Time min:s |
|---|---|---|---|
| 1 | 1 | Histoclear | 3:00 |
| 2 | 2 | Histoclear | 3:00 |
| 3 | 3 | Histoclear | 5:00 |
| 4 | 4 | Absolute Ethanol | 1:00 |
| 5 | 5 | Absolute Ethanol | 1:00 |
| 6 | 6 | 95% Ethanol | 1:00 |
| 7 | Wash5 | Water | 1:00 |
| 8 | 9 | Hematoxylin | 3:30 |
| 9 | Wash4 | Water | 2:00 |
| 10 | 7 | .1% HCl in 70% Ethanol | 0:01 |
| 11 | Wash3 | Water | 1:00 |
| 12 | 10 | Scotts Water | 3:30 |
| 13 | Wash2 | Water | 1:00 |
| 14 | 11 | Eosin | 0:30 |
| 15 | 12 | 95% Ethanol | 1:00 |
| 16 | 14 | 95% Ethanol | 2:00 |
| 17 | 15 | Absolute Ethanol | 1:00 |
| 18 | 16 | Absolute Ethanol | 1:00 |
| 19 | 17 | Histoclear | 1:00 |
| 20 | 18 | Histoclear | 1:00 |
| 21 | Exit | Histoclear | - |

**Supplemental Table 1.** The H&E automated staining protocol with steps, reagents and slide times shown.
